# Gonadal and sex chromosomal contributions to sex differences in mammalian brain organization

**DOI:** 10.64898/2026.01.10.698782

**Authors:** Jason P. Lerch, Juliana Shin, Jacob Ellegood, Haley Hrncir, Brian J. Nieman, Allan Mackenzie-Graham, Xia Yang, Arthur P. Arnold, Armin Raznahan

## Abstract

Sex differences in brain development may contribute to well-known sex differences in behavior and neuropsychiatric risk - making it important to comprehensively map and mechanistically annotate normative sex differences in brain organization. Most sex differences in mammalian brain organization have previously been attributed to differential effects of gonadal hormones based on outcomes from rodent endocrine manipulations at canonical subcortical foci of male-biased brain volume. However, a systematic quantification of both gonadal and sex chromosome dosage (SCD) contributions across all regions of sex-biased brain volume has been lacking. Here, using structural neuroimaging scans from wild-type (n=670) and transgenic mice that dissociate gonadal, X-, and Y-chromosome effects (n=181), we show that: many more brain regions are volumetrically sex-biased than previously recognized; gonadal effects dominate throughout this expanded map of sex differences; several regions also show prominent SCD contributions to anatomical sex differences, which can both reinforce or counteract gonadal effects in a regionally specific manner. Targeted single nucleus RNA sequencing at a region of female-biased cerebellar volume reveals that combined gonadal and SCD effects also drive sex-biased cellular gene expression. These findings revise our understanding of the spatial distribution and causal basis of sex-differences in the mammalian brain, illuminating a key axis of biological variation in health and disease.

## Introduction

Identification and mechanistic study of sex-differences in the brain is important for understanding sex-biased behavior in health and illness^1^. Given limited experimental access to human brain tissue, much of this knowledge comes from rodents, where histological work has identified reproducible foci of sex differences in cell count and volume - larger in males for the bed nucleus of the stria terminalis, medial amygdala and sexually dimorphic nucleus of the medial preoptic areas (BST, MEA, and SDN-MPO), and larger in females for the Anteroventral Periventricular Nucleus (AVPV)^2–5^. These findings established a long-standing “gonad-centric” model of sex-biased brain development - rooted in decades’ of research on gonadal regulation of rodent brain development^6^, and solid experimental evidence linking selected foci of sex-biased anatomy to gonadally determined sex differences in circulating testosterone^7,8^.

More recent access to transgenic mice has suggested that sex differences in the brain may not be purely of gonadal origin, but also shaped by another core biological source for phenotypic sex differences that is shared by all placental mammals: the XX vs. XY difference in sex chromosome complement formed by disparate X- and Y-chromosome dosage (henceforth sex chromosome dosage, SCD)^9–11^. In particular, high throughout murine structural magnetic resonance imaging^12^ (sMRI-which can successfully recover and extend upon classical foci of sex-biased anatomy from histological research^13,14^) has revealed that regional volume of the mouse brain is also sensitive to variations in SCD^15–18^. These observations suggest that sex-biased brain anatomy may arise through combinatorial effects of gonads, X-chromosome dosage (XCD), and Y-chromosome dosage (YCD), but no brain-wide test of this hypothesis exists - despite its potential to revise classical models^19^.

Here, we apply new analytic procedures in expanded sMRI datasets (total n=851) to first provide an unprecedentedly fine-grained (40 µm isotropic brain-wide) map of sex-biased brain volume in wild type mice (*MouseBase*, n=670), and then systematically annotate this map according to the relative effects of gonadal type (testes vs. ovaries), sex chromosome complement (XX vs. XY), XCD, and YCD as estimated using the Sex Chromosome Trisomy (SCT^20^) mouse model (n=181). Guided by these findings we focus on a newly identified region of female-biased cerebellar volume and perform sMRI-targeted single-nucleus RNA sequencing (snRNAseq) in SCT mice (n=24) to test for combined gonadal and chromosomal contributions at underlying cellular and molecular levels of brain organization.

We find that over 50% of the mouse brain shows statistically significant volumetric sex differences by sMRI with over 10% showing moderate to strong effect sizes. Contrary to the traditional emphasis on foci of male-biased volume in the mouse brain, the majority of sex-biased regions are female-biased. We replicate the classical finding of major gonadal contributions at canonical foci of both male- and female-biased brain volume, and further show that these dominant gonadal influences also operate throughout the substantially expanded map of sex-biased brain anatomy we report herein. Moreover, we also find a significant contribution of sex chromosomal effects to sex differences in regional brain anatomy - which are spatially coordinated with gonadal effects and reflective of the balance between volumetric influences of XCD and YCD. We provide a spatially fine-grained mechanistic annotation of all sex-biased regions, which reveals how sex chromosomal effects sometimes add to and sometimes counter the volumetric effects of gonadal type. Targeted snRNAseq within a region of female-biased cerebellar cortex shows that gonadal and sex chromosomal effects also jointly shape cell-type–specific sex differences in brain gene expression - helping to clarify the cellular and molecular basis for observed volume differences.

Taken together, these results: provide a new reference map of sex-biased murine brain organization (https://osf.io/4cbas); reveal coordinated gonadal–chromosomal influences on anatomy and gene expression; and, offer a framework for parsing sex differences in the human brain, where some foci of sex-biased brain volume are shared with mice^14^.

## Results

### A reference map of sex differences in murine brain volume

We collated a large sMRI dataset of young adult wild-type mice (*MouseBase*, n=670, 336 male, mean age of P60, age range of P56 to P112, **Methods**) gathered at the Mouse Imaging Centre (MICe, Toronto, Canada) as controls for a neurodevelopmental study^21^. By applying a well-established deformation-based morphometry pipeline^22^ to *MouseBase*, followed by voxel-wise linear mixed-effects models, we estimated the effect of sex on volume at each of ∼7.5M 40 µm isotropic voxels across the entire mouse brain (**Methods**). This procedure enabled us to quantify the proportion of brain voxels showing volumetric sex differences across significance and effect size thresholds (**Fig 1 a-c**).

**Figure 1.**
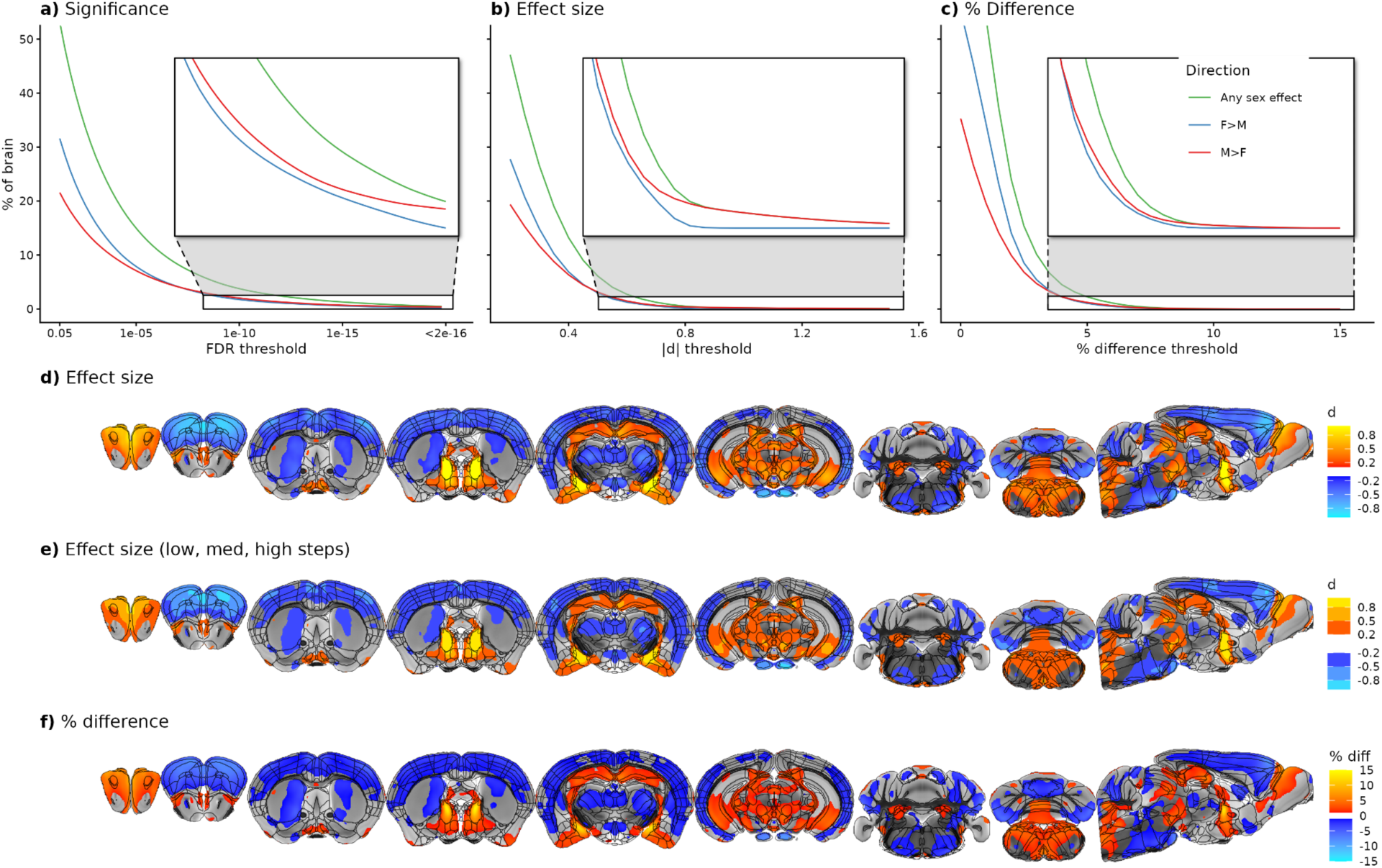
Statistical significance, magnitude and spatial distribution of volumetric sex differences in the mouse brain. **a, b, c.** The proportion of all ∼7.5M 40µm^3^ voxels in the mouse brain showing a sex difference in volume (any direction - green; female-biased - blue; male-biased - red) at varying thresholds for statistical significance (**a**), effect size (**b**) and percent volume difference (**c**). **d, e, f**. Voxel-wise maps for selected slices of the mouse brain showing statistically significant sex-biased volume (FDR q<0.05) visualized as (**d**) continuous effect size, (**e**) effect size in steps of low (0.5>|d|>0.2), medium (0.8>|d|>0.5), and high (|d|>0.8) effects, and (**f**) percent volume difference. Additional slices of the brain map in d) are provided in **Fig S1**. Colors encode the direction and magnitude of the observed volumetric sex difference (female> male - blue; male>female - red). Regions falling within effect size steps of e) are listed in **Table 1**.

After accounting for whole-brain volume, over half (53%) of all brain voxels showed a statistically significant sex difference in volume after False Discovery Rate (FDR^23^) correction for multiple comparisons across voxels with q (the expected proportion of falsely rejected null hypotheses) set at 0.05. Slightly more of these voxels (59%) were female biased (i.e. larger mean volume in females vs. males) than male biased (**Fig 1a**). Collectively, these voxels showed a wide variation in the magnitude of observed sex differences as estimated by both standardized effect sizes (0.17 < |d| < 2.3, **Fig 1b**) and percent differences [0.99 < ((|M-F|/F)*100) < 15.2, **Fig 1c**]. Voxels showing the very largest magnitude sex-differences within this set (i.e. |d|>0.98, **Figs 1b** inset) were all male-biased.

Large effect size (|d|>0.8) volumetric sex difference by sMRI encompassed all classical foci of male-biased brain volume described by prior histological research in rodents^8,24–26^ (**Fig 1e**, **Table 1, Fig S1**) - medial preoptic area of the hypothalamus (MPO), bed nucleus of the stria terminalis (posterior division, pBST) and medial amygdalar nucleus (posterodorsal part, pdMEA) - as well as histologically undescribed regions of sex-biased volume in the main olfactory bulb (male-biased), as well as prelimbic cortex, secondary motor cortex and pontine gray (all female-biased).

**Table 1.**
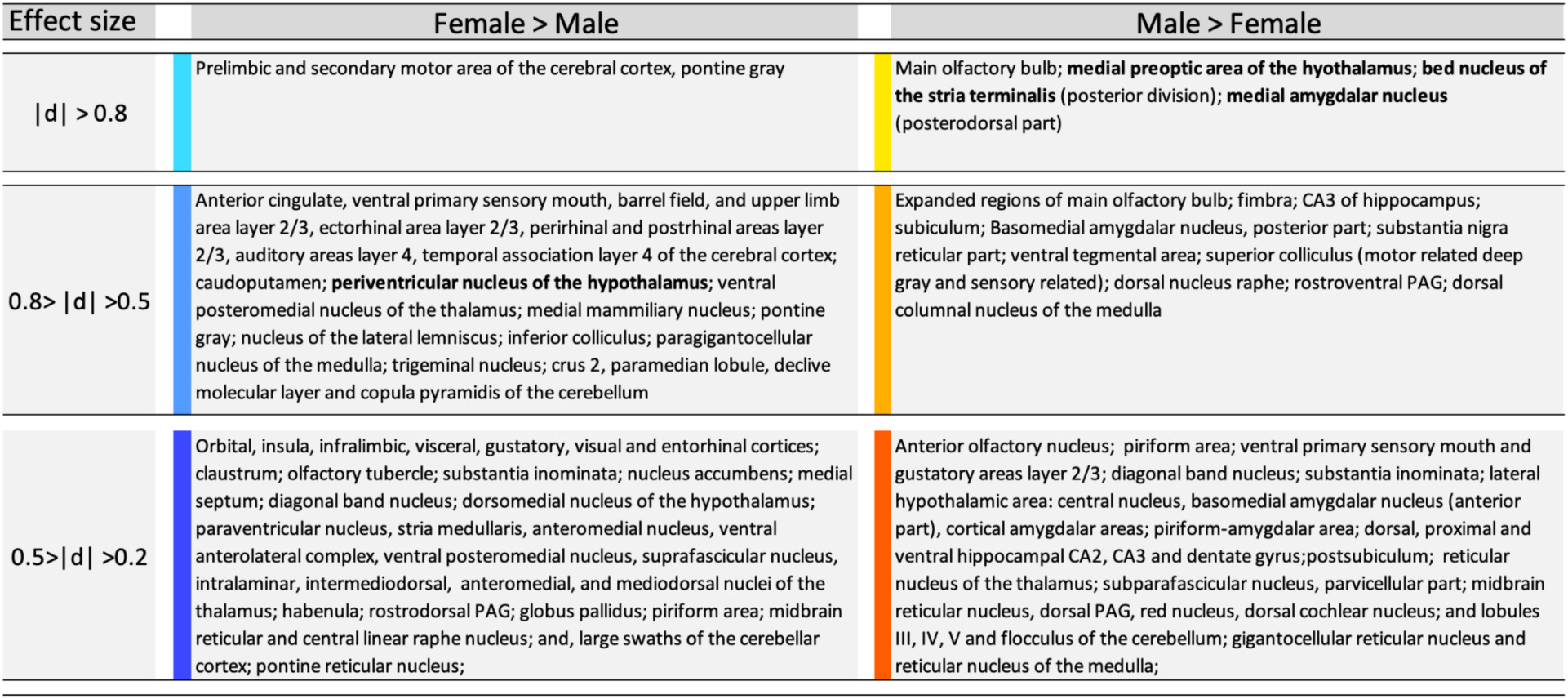
Sex-biased regions in the mouse brain from Fig 1, stratified by direction of sex-bias (columns) and effect size (rows). Color codes match **Fig 1e**. Bold text denotes regions of volumetric sex differences in the mouse brain defined by prior histological research.

At a moderate effect size threshold (|d| > 0.5) there were expanded territories of sex-biased volume around the aforementioned high effect size foci, as well as several additional regions of sex-biased volume which are all largely undescribed by prior histological studies (**Fig 1e**, **Table 1, Fig S1**). Significantly male-biased regions at this effect size threshold included additional parts of the amygdala, hippocampal CA1, numerous motor- and behavioral-state related midbrain nuclei and the dorsal medulla. Significantly female-biased regions at this effect size threshold included: the AVPV nucleus of the hypothalamus; parahippocampal regions and the mammillary nuclei to which they project; and numerous components of somatosensory and auditory systems in the cortex, thalamus, midbrain, pons and cerebellum.

At a small effect size threshold (|d| > 0.2) the spatial extent of these novel sex-biased foci expanded further still (**Fig 1e**, **Table 1, Fig S1**) to encompass numerous additional foci of male-biased (expanded amygdalo-hippocampal regions; olfactory regions of the basal forebrain; lateral hypothalamus; motor- and sensory related components of the midbrain and medulla; and subregions of the cerebellum) and female-biased (claustrum; expanded regions of limbic cortex, thalamus, and cerebellum; striatum, periaqueductal gray and midbrain reticular nuclei) volume.

In gestalt (**Fig 1d-f, Fig S1**), volumetric sex differences appear to follow a broad anatomical patterning across the mouse brain: volume tends to be male-biased throughout the olfactory system, extended amygdala, hippocampus, and components of the extrapyramidal motor system, but female-biased throughout much of the cerebral cortex, striatum, thalamus, and cerebellum.

Thus, murine sMRI recovers all classical histologically described foci of sex-biased volume in the mouse brain - buttressing our observation of widespread volumetric sex differences outside these classically reported foci. These additional foci of sex-biased volume: include medium-large effect sizes; are widely distributed across multiple compartments of the murine brain; and, encompass both male- and female-biased regions that apparently concentrate within distinct functional systems of the murine brain. To facilitate use of this new information by the wider research community, we share all key voxel-wise maps of sex differences in murine brain volume (FDR q value, effect size, and percent difference), a co-registered parcellation of the mouse brain, and a corresponding anatomical space to which these statistical and parcellation files are registered (https://osf.io/4cbas).

### Spatial covariation between volumetric sex differences and the volumetric effects of gonads and sex chromosomes

We next sought to annotate sex-biased regions in the mouse brain according to the causal role of gonads and sex chromosomes - the two core biological bases for phenotypic sex differences in all placental mammals. For this purpose, we first specified a “sex-biased mask” (SBM) for mechanistic annotation - comprising all voxels showing a statistically significant volumetric sex difference (FDR q<0.05) in the *MouseBase* dataset of a |d| ≽ 0.3 effect size. Regional estimates of sex effects on volume based on a previously described parcellation of the mouse brain^27^ are provided as a table at https://osf.io/4cbas.

Before annotating SBM voxels defined in *MouseBase* for gonadal and chromosomal effects estimated by sMRI in SCT mice (see below), we confirmed strong cross-voxel concordance in volumetric sex differences (r=0.61, **Fig. S2a,b**, **Methods**) between *MouseBase* (wild type males vs. females) and the SCT model (XY-testes vs. XX-ovaries groups) despite the different genetic backgrounds between *MouseBase* and SCT cohorts (predominantly C57BL/6 vs. outbred MF1). This concordance justified modeling gonadal and sex chromosome effects in SCT sMRI data, and then using these effects to annotate normative sex effects described in the larger *MouseBase* dataset to which SCT data had been spatially coregistered (**Methods**). Through this process, we achieve a one-shot integrated description and mechanistic dissection of sex-biased murine brain volume at 40 µm isotropic resolution.

The SCT mouse model is previously described^20^. Briefly, breeding XY^-^(*Sry*^+^) male mice (which have the testes-determining *Sry* gene relocated from the Y-chromosome to an autosome - producing X and Y^-^ sperm each with or without autosomal *Sry*) with XXY^-^ females (producing X and XY^−^ eggs) generated all 8 groups of the SCT mouse model consisting of four sex chromosome karyotypes - XX, XY, XYY or XXY - each with either ovaries or testes (**Fig 2a**). We used two linear models in the SCT dataset (**Methods**) to estimate the voxelwise influences of gonads and sex chromosomes on murine brain volume: one in the “Four Core Genotypes” (FCG)^9^ subset of the SCT model (XX-ovaries, XX-testes, XY-ovaries and XY-testes, combined n=97) to separately estimate voxel-wise effects of gonad type (testes vs. ovaries, TvO) and sex chromosome complement (XYvXX); and another in all eight SCT groups (n=181) to separately estimate voxel-wise effects of increasing X-chromosome dosage (XCD: 1 or 2) and Y-chromosome dosage (YCD: linear effect of increasing dosage across the range 0, 1, 2) while accounting for gonadal type (TvO). **Fig 2b** reproduces the effect size map for volumetric sex differences in the *MouseBase* SBM, alongside effect size maps for the following four gonadal and sex chromosome terms estimated from models within the SCT dataset: TvO (as estimated from the full SCT cohort of 8 groups rather than the FCG subset of 4 groups), XYvXX, XCD, and YCD.

**Figure 2.**
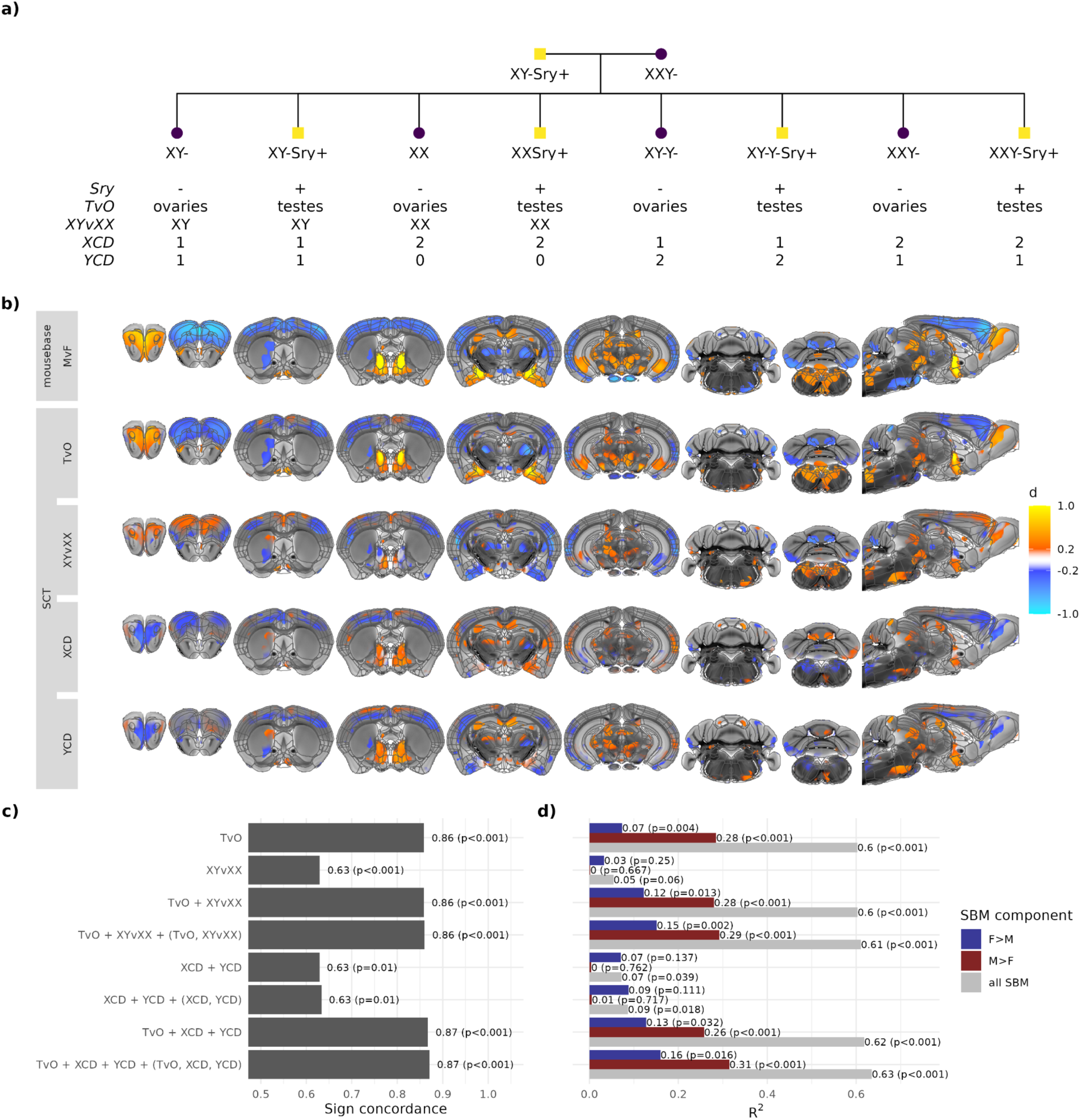
Estimating gonadal and sex chromosome effects on regional brain volume and their spatial congruence with volumetric sex differences. **a,** Schematic of the cross producing all eight groups of the SCT model, with details of the gonadal type (Testes vs. Ovaries, TvO), sex chromosome complement (XYvXX), and X- and Y-chromosome dosage (XCD, YCD, respectively) of each group. **b** Voxel maps for selected coronal and sagittal views of the mouse brain showing the effect size of normative volumetric sex differences from *MouseBase* SBM (from Fig 1), as well as volumetric effects of gonadal type (TvO), sex chromosome complement (XXvXY), XCD, and YCD as computed within the SCT mouse cohort and thresholded at |d| ≥ 0.2. **c,d** Bar charts for diverse cross-voxel models predicting spatial variation in volumetric sex differences within the *MouseBase* sex-biased mask (SBM) as a function of spatial variation in the voxel-wise effects of terms in the SCT model (blue - female-biased SBM (F>M); red - male-biased SBM (M>F); gray - whole SBM). **c** shows the proportion of SBM voxels with sign concordance between the direction of sex-biased volume predicted by each linear model and the direction of observed volumetric sex differences in *MouseBase*. **d** shows the proportion of spatial variation in observed volumetric sex differences within the *MouseBase* SBM that is explained by spatial variation in the effects of different combinations of SCT terms.

Qualitative comparison of the effect size maps in **Fig 2b** revealed that although normative male vs female (MvF) sex differences in the SBM most closely align with gonadal effects (general concordance between MvF and TvO with notable exceptions in the dorsal hippocampus, superior colliculus, visual, auditory and entorhinal cortices), there are also several regions where observed sex differences are also congruent with sex chromosome effects (primary sensory cortices, ventral caudoputamen, dorsal hippocampus, ventral thalamic nuclei, superior colliculus, cerebellar cortex, cerebellar nuclei, and medulla). Supporting these qualitative impressions, cross-voxel spatial Pearson correlations within the SBM between the MvF effect size from *MouseBase* and each of the SCT term effect sizes were as follows: TvO 0.78; XYvXX 0.24; XCD 0.02; YCD 0.18 (**Fig S2c**). We also observed qualitative spatial alignments amongst TvO, XYvXX, XCD and YCD effect maps within the SBM (**Fig 2b**), which were confirmed by quantifying cross-voxel correlations amongst these terms (**Fig S2c**). These correlations were strongest between XCD and YCD (r=0.7), but also substantial between each of these terms and XYvXX (r= −0.42, and r=0.22, respectively), as well as between XYvXX and TvO (r=0.34).

Given the strong spatial alignment between XCD and YCD effects on brain volume, we derived maps for their algebraic sum (YCD + XCD, **Fig S2d top row**) and difference (YCD-XCD, **Fig S2d bottom row**). The (XCD+YCD) map identified regions whose volume scaled with total sex chromosome count, some overlapping normative sex differences (e.g. BST, MEA, limbic cortex), although the (XCD+YCD) map showed a weak global spatial correlation with sex effects (r=0.12, **Fig S2c**). The (YCD-XCD) map was more strongly correlated with normative sex differences (r=0.23, **Fig S2c**) and at a similar magnitude to the correlation between XYvXX and sex effects (r=0.24, **Fig S2c**) - reflecting the strong correlation between (YCD-XCD) and XYvXX effects (r=0.8, **Fig S2c**). Thus, the disparity between YCD and XCD effects as estimated in the context of aneuploidy can strongly predict the volumetric effects of normative sex differences in chromosome dosage between XY males and XX females.

To quantify the relative contributions of gonadal and sex chromosome effects to anatomical sex differences, we used spline models (**Methods**) to relate inter-voxel variation in *MouseBase* sex differences (**Fig. 1d, Fig. S1, Fig. 2b**) to combinations of TvO, XYvXX, XCD, and YCD effects estimated in the SCT model (**Fig. 2b**). These models were fit using SBM sex differences calculated in half of the *MouseBase* dataset, and model performance was evaluated against SBM sex difference in the remaining half using 10-fold cross-validation (**Methods**). For each model, we calculated: (i) the proportion of SBM voxels where predicted sex differences matched the direction of observed sex differences (**Fig. 2c**); and (ii) the magnitude of spatial variance explained in *MouseBase* sex differences when considering all SBM voxels as well as their male- and female-biased subsets (**Fig. 2d**, gray, blue and red bars, respectively). Results were compared against null models generated from 100 random permutations of sex for the test dataset within the split half scheme described above (**Methods**).

These analyses revealed that the direction of volumetric sex-difference could be correctly predicted for >85% of SBM voxels using gonadal effects alone (p<0.001), but only >60% using either XYvXX or combined XCD/YCD terms in isolation (p<0.001 and .01, respectively) (**Fig 2c**). Analyses of variance explained (**Fig. 2d**) confirmed a close alignment between *MouseBase* sex differences and TvO effects across the whole SBM (R²>0.6, p<0.001), and more strongly in male-vs. female biased SBM subsets (R²=0.28 vs. 0.08, both p<0.001). In contrast, sex chromosome terms alone achieved a much weaker prediction of regional volumetric sex differences across the SBM as a whole (maximal R² =0.09, p<0.001). However, inclusion of sex chromosome terms significantly boosted prediction of regional sex differences within female-biased parts of the SBM compared to a model containing gonadal terms alone (**Fig. S3a**). For example, compared to a model with a sole gonadal term (TvO), a model incorporating additional effects of sex chromosome complement and their interaction with gonadal effects [i.e. MvF ∼ TvO + XYvXX + (TvO, XYvXX)] predicted significantly more variance in the spatial patterning of regional sex differences within female-biased regions of the SBM (R² improvement=0.08, p=0.02).

Taken together, these findings establish that regional sex differences in brain volume closely track gonadal effects on regional brain volume across the expanded SBM we newly describe in *MouseBase* - which extends far beyond the classical foci of sex-biased volume with experimental evidence for gonadal effects from rodent endocrine manipulations. We also show that the spatial alignment between gonadal effects and sex differences is stronger in male- vs. female-biased regions of the SBM. However, despite the dominance of gonadal effects over sex chromosomal effects in prediction of regional sex differences, we find that prediction of regional sex differences within female-biased brain regions benefits from combined consideration of both gonadal and chromosomal effects. By disambiguating effects of gonadal type and sex chromosome complement on brain volume, we also discover a hitherto unappreciated spatial coordination between these two effects. Sex chromosome complement effects are in turn predicted by the differences between XCD and YCD effects on anatomy, although these differences are nested within a general spatial concordance between effects of these two heteromorphic chromosomes on regional brain volume.

### Spatially fine-grained annotation of volumetric sex differences for the combined contributions of gonadal and chromosomal effects

The analyses above quantify the overall spatial correlation of gonadal and sex chromosome effects with sex-differences in regional brain volume (**Fig 2, Fig S2, S3**) but not the relative influence of gonadal and chromosomal effects at each sex-biased focus. To resolve this, we used the aforementioned bivariate spline model for prediction of regional sex differences as a function of both gonadal type and sex chromosome complement [(MvF ∼ s(TvO) + s(XYvXX) + s(TvO, XYvXX), **Fig. 2**, **Methods**)]. This model allowed plotting sex-difference magnitudes across combinations of TvO and XYvXX effects (**Fig. 3a,b**) to reveal that: (i) the largest sex differences (i.e. regions where predicted sex differences are |d|≥0.5, as bounded by green lines in **Fig. 3a**) occur where strong gonadal effects coincide with opposing chromosome effects (e.g., M>F where T>O but XX>XY); but (ii) most sex-biased voxels (larger dots in **Fig. 3b**) reflect concordant gonadal and chromosomal effects (e.g., M>F where T>O and XY>XX).

**Figure 3.**
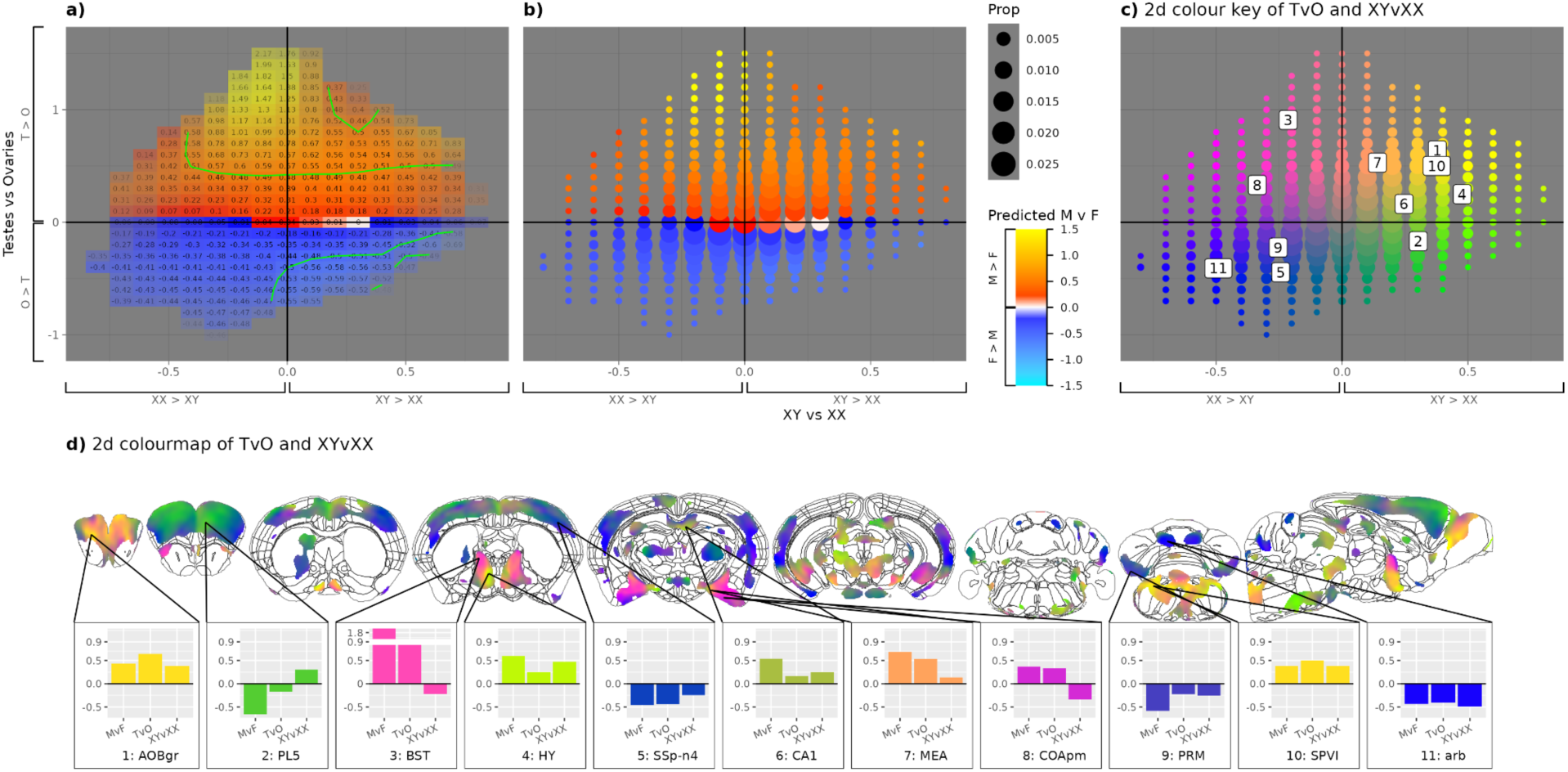
Voxelwise mechanistic annotation of sex differences in murine brain volume. **a,** Raster plot showing the predicted effect size of volumetric sex differences (red - male-biased; blue - female-biased, white - near zero) based on a cross-voxel estimation of observed volumetric sex differences within the *MouseBase* sex biased mask (SBM) from observed volumetric effects of gonadal type (Testes vs. Ovaries; TvO) and sex chromosome complement (XXvXY) within the SCT dataset (**Methods**). Estimated sex differences are shown for each combination of TvO (y-axis) and XXvXY (x-axis) effect sizes. Green contour lines demarcate predicted sex differences with an absolute effect size > 0.5 (i.e between upper two green lines and below lowest green line in a). **b,** A plot with the same information as panel a, but use of dot size to reflect the proportion of sex-biased voxels at each combination of effect sizes for gonadal type and sex chromosome complement. **c,** A plot with the same axes and dot size information as panel b, where dot color is parameterized according to a color space that captures graded variations in the relative effect size of gonadal type and sex chromosome complement. Inset labels show the location of selected brain regions. **d,** Selected coronal and sagittal slices of the mouse brain showing sex-biased voxels of the MouseBase SBM color coded by the colorspace from panel c - providing a mechanistic annotation of normative sex differences in murine brain volume for the relative magnitude and direction of volumetric influences from gonadal type and sex chromosome complement. Labels specify brain regions that illustrate the heterogeneous mechanistic contributions to volumetric sex differences. Underlying bar plots show observed volumetric effects of sex (normative sex differences from MouseBase), gonadal type and sex chromosome complement (both from the SCT cohort) for selected brain regions highlighted in panels c and d. **Abbreviations.** AOBg - accessory olfactory bulb, granular layer; PL5 - prelimbic area, layer 5; BST - bed nucleus of the stria terminalis; HY - hypothalamus; SSp-n4 - primary somatosensory area, nose, layer 4; CA1 - hippocampal field CA1; MEA - medial amygdalar nucleus; COApm - cortical amygdalar area, posterior part; PRM - paramedian lobule of the cerebellum; SPVI - spinal nucleus of the trigeminal, interpolar part; arb - arbor vitae of the cerebellum.

Creating a color-space to encode all TvO–XYvXX combinations (**Fig. 3c**) and projecting these onto SBM voxels (**Fig. 3d**), provided a fine-grained map of gonadal vs. chromosomal contributions to sex differences that revealed marked spatial heterogeneity. Male-biased volume reflected either: (i) summative, moderate, concordant effects of gonadal type and sex chromosome complement (T>O and XY>XX; yellow–khaki in **Fig. 3d,e**) in accessory olfactory bulb (AOBgr), hypothalamus, hippocampal CA1, MEA, and the spinal nucleus of the trigeminal nerve (SPVl) (Regions 1,4,6,7,10, respectively); or (ii) strong gonadal effects outweighing opposing sex chromosome complement effects (T>O and XX>XY; magenta–purple in **Fig. 3d,e**) in BST and the cortical amygdala (COApm) (Regions 3,8, respectively). A smooth transition between these mechanisms was observed from hypothalamus to BST. Female-biased regions showed similar mechanistic diversity. Some reflected concordant effects of gonadal type and sex chromosome complement (O>T and XX>XY; blue in **Fig. 3d,e**), such as in the primary somatosensory cortex (SSp), cerebellar paramedian lobule (PRM), and cerebellar white matter (arb) (Regions 5,9,11, respectively). Others showed counterbalanced contributions (O>T and XY>XX; green in **Fig. 3d,e**), such as in the prelimbic cortex (PL5, Region 2). A graded shift between these mechanistic paths was evident across the frontal cortex, from ventrolateral insula to dorsomedial prelimbic regions. Voxel-wise model comparisons (**Methods**) revealed that consideration of sex chromosome effects significantly improves prediction of regional volumetric sex differences above and beyond gonadal effects alone in regions of both male-biased (BNST, medial amygdala, hippocampus) and female-biased (prelimbic, secondary motor, entorhinal, and cerebellar cortices) brain volume (**Fig S3b**).

Thus, regional sex differences in brain volume are underpinned by spatially graded mechanisms reflecting varying combinations of gonadal and chromosomal contributions. These generally operate in concert and congruently with the observed direction of sex-biased anatomy. However, in regions with the largest effect size sex differences, although gonadal influences remain congruent with the observed volumetric sex difference, chromosomal influences run in the opposite direction, as if to partly counter gonadal effects.

### Combined contributions of gonadal and chromosomal effects to sex differences at single cell resolution

Observing that sex-biased brain volume reflects the combined effects of gonadal and sex-chromosomal factors (**Figs 2 and 3**) suggests that similar combinatorial effects may also operate at underlying cellular levels of brain organization. We tested this hypothesis through single nucleus RNAseq (snRNAseq) analysis of cellular proportions and cell-type specific gene expression in SCT mice - targeting a prominent and previously undescribed region of female-biased cerebellar volume exhibiting mixed gonadal and chromosomal effects in our sMRI analysis: the paramedian lobule of the cerebellum (PRM, **Fig 1**, **Table S1**, **Fig 3, Structure 9 PRM**).

We generated snRNAseq libraries (using 10x Chromium v3.1, **Methods**) from PRM samples of 24 mice encompassing individuals from each of the 8 SCT model groups (mean age 65 days, range 60-70, no difference in age by genotype F=0.7, p=0.7). After QC (**Methods)**, 194,720 cells remained, clustered into 14 canonical cerebellar cell types based on a reference study^28^ (**Fig. 4a**; **Methods**), each represented across all SCT groups (**Fig. S4a**). Cellular proportions did not vary across SCT groups (**Fig 4b**, Omnibus F test for group*cell-type interaction, F=0.6, p=1) - implying that the observed effects of sex, gonads and chromosomes on PRM volume (**Fig 3**) are more likely to reflect variations in cell density or shape than cell type composition.

**Figure 4.**
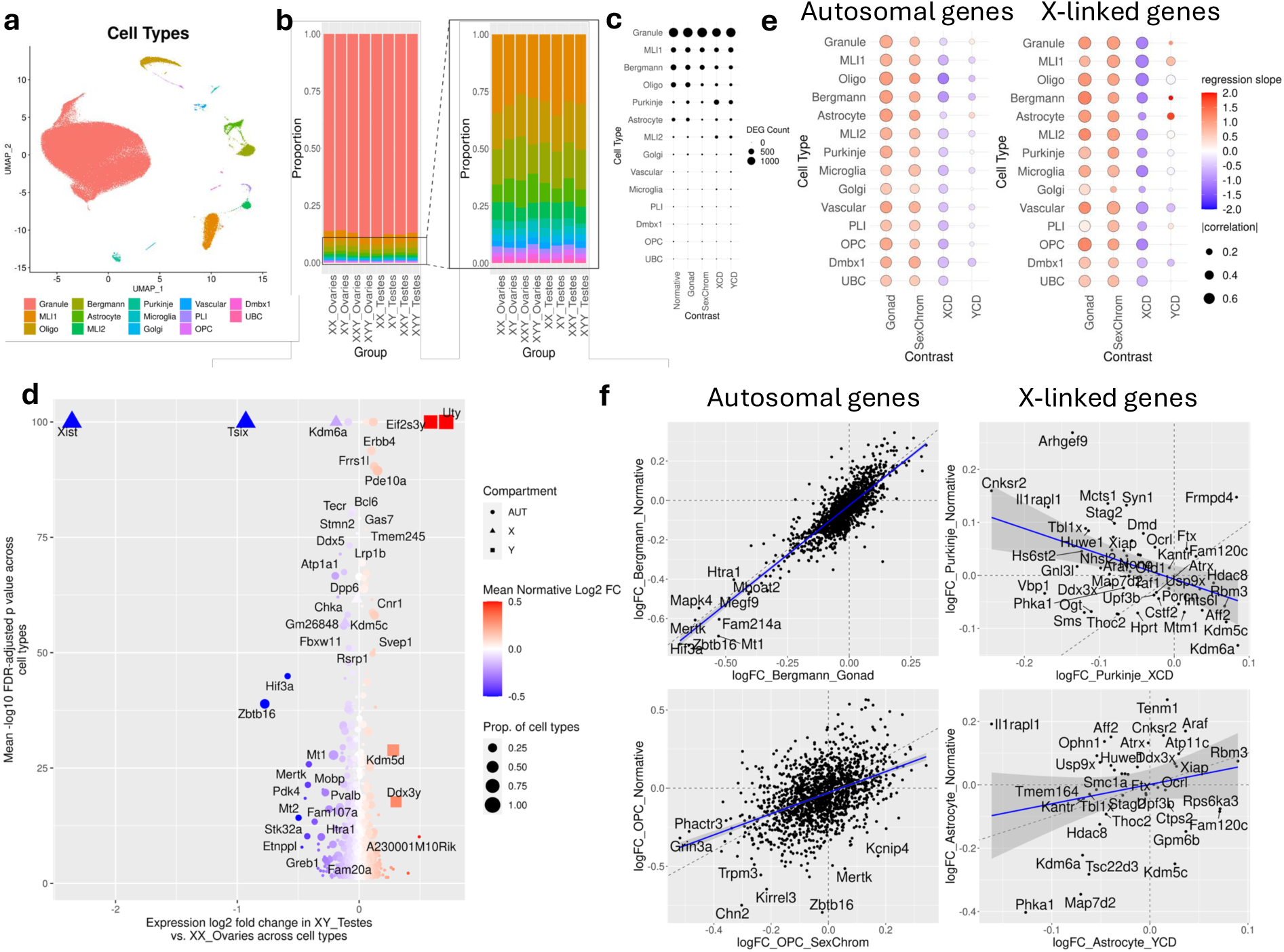
Transcriptomic analysis of sex, gonadal and sex chromosome dosage effects on the cerebellum at single cell resolution. **a,** UMAP showing clustering of cerebellar cell types. **b**, Barplot showing cell type proportions in all 8 groups of the SCT mouse model, shown for all cell types (left) and excluding Granule cells which account for the vast majority of cells in all groups (right). **c**, Dotplot showing number of statistically significantly differentially expressed genes (FDR q<0.05, DEGs) for each of 5 observed contrasts (columns) in each of 14 observed cell types. **d,** Scatterplot showing 2142 genes which are normative sex DEGs in at least one cell type. Dot size encodes the proportion of all 14 cell types that the gene is a DEG in: dot shape encodes whether gene is autosomal, X-linked or Y-linked; x-axis shows mean log2 expression fold change across cell types (red - M>F; blue - F>M); y-axis shows mean -log10 p value for significance of normative sex difference across cell types. **e,** Dotplots showing the similarity (dot size: fold-change correlation across genes, color saturation: fold-change regression slope) between normative sex differences in gene expression and expression effects of 5 different contrasts from the SCT model (columns) for each cell type (rows) - shown separately for autosomal (left) and X-chromosome genes (right) that showed a significant normative sex differences in expression in at least one cell type. **f,** Illustrative fold-change correlation scatterplot for cell types from **e** - shown separately for autosomal genes (left column = in Bergmann and OPC cells) and X-chromosome genes (right column - in Purkinje neurons and Astrocytes). The x=y identity line is dotted gray and the regression fit line is solid blue.

Using the limma R package^29^ (**Methods**), and identical linear models to those applied in neuroimaging analysis, we profiled transcriptomic effects of sex (i.e. expression differences between XY-testes vs. XX-ovaries SCT groups), gonads (TvO), sex chromosome complement (XYvXX), XCD, and YCD on each cerebellar cell type (Gene-level results for all terms available at: https://osf.io/4cbas). Ranking of cell types by the number of differentially expressed genes (DEGs at FDR q<0.05) was broadly consistent across all 5 of these estimated effects (**Fig 4c**). Sensitivity analyses revealed that this recurrent cell type ranking by DEG count tracked with cell type abundance (**Fig S4b**).

Sex was associated with 2132 unique DEGs across cell types. The most recurrent sex-related DEGs across cell types - which often also showed greatest mean fold change and statistical significance across cell types (**Fig 4d**) - were (i) *Xist* and *Tsix* (F>M expression) - the pair of antisense lncRNA transcripts that orchestrate X-chromosome inactivation (XCI^30,31^) in females, and (ii) *Uty*, *Ei2s3y*, *Kdm5d* and *Ddx3y* (M>F expression) - four Y-linked gametologue genes previously reported as being highly dosage sensitive in both bulk^32,33^ and single cell^34^ transcriptomic studies of the mouse brain. The majority of sex DEGs (n=2076) were autosomal, although these showed substantially less recurrence across cell types and smaller mean expression fold changes than the previously mentioned sex-chromosome genes (**Fig 4d**). Supplementary analyses confirmed more inter-cell-type consistency in sex effects on expression of sex chromosome genes than autosomal genes (paired t-test: t=3.3, p=0.002, **Fig S4c**). Gene Ontology (GO) enrichment of sex DEGs per cell type (**Fig. S4d,e**, **Table S2**) implicated broad neuronal and glial processes (e.g. synaptic signaling, gliogenesis, cerebellar morphogenesis, cytoskeleton), alongside cell-type specific pathways (e.g. extracellular matrix in Bergmann cells, immune response in microglia). Thus, sex differences in gene expression appear to impact diverse aspects of biological structure and function in the cerebellar PRM, including several that could plausibly contribute to the sex-biased volume of this region observed by sMRI (**Fig 1**, e.g. via sex differences in neurite anatomy, synapse size/density or the extracellular matrix).

We next used cross-gene fold change correlation and regression analyses (**Fig 4e, Methods**) to test if and how normative sex differences in gene expression are related to gonadal and sex chromosomal effects (**Methods**). For almost all cerebellar cell types, normative sex differences in autosomal gene expression (MvF) were positively correlated with the expression effects of both gonadal type (TvO) and sex chromosome complement (XYvXX) (**Fig 4e left,** for n=2079 autosomal sex DEGs in at least one cell type) - indicating a contribution from both factors in shaping normative sex differences in autosomal gene expression. Strikingly, this same phenomenon was also observed for sex differences in X-chromosome gene expression (**Fig 4e right,** for n=57 sex DEGs in at least one cell type - excluding Xist, Tsix and all 5 Y-linked genes in our dataset due to their outlier expression changes) - suggesting a rarely discussed capacity of gonadal effects to shape sex differences in X-chromosome gene expression (in addition to the expected influence of sex differences in X-chromosome count). For both autosomal and X-linked genes, and across most cell types, sex differences in expression were consistently aligned with XCD effects (i.e. negative regression coefficients, **Fig. 4e**), while correlations between sex and YCD effects on expression were weaker and more cell-type specific (**Fig. 4e**). **Fig. 4f** provides scatterplots illustrating correlations between sex differences in gene expression and expression effects of selected sex-related factors for selected cell types.

Despite the distinct relationships of XCD and YCD effects on gene expression with sex differences in gene expression (**Fig 4e**), we found that XCD and YCD effects on gene expression were positively correlated with one another for most cell types (**Fig. S4f**). This combination of findings suggests that the regulatory mechanisms driving alignment between XCD and sex effects on expression are non-overlapping with those driving alignment between XCD and YCD effects. Moreover, the consistent alignment we observe between XCD and YCD effects on gene expression echoes the tight correlation between XCD and YCD effects on brain anatomy (**Fig 2, S2**).

Taken together, these transcriptomic findings suggest that sex, gonadal and sex chromosomal effects on brain volume may not necessarily involve sex differences in cell type composition, but could instead potentially reflect the influence of these factors on aspects of cellular morphology. Moreover, it appears that the combinatorial influence of gonadal and chromosomal effects on sex-biased anatomy (**Fig 2,3**) is accompanied by similar combinatorial effects of these factors in sex-biased gene expression (**Fig 4**). Finally, both brain anatomy and cellular gene expression show highly convergent effects of XCD and YCD. This recurrent theme of convergent effects is surprising given the dramatic heteromorphism of X- and Y-chromosomes - and points to a conserved causal role for one or more of the small handful of genes that these two chromosomes still share.

## Discussion

Our study advances understanding of the spatial distribution and mechanistic bases for neuroanatomical sex differences in mice, a dominant model for mammalian brain development. We review each advance in turn, before considering implications for other mammals, including humans.

### Updating the anatomical distribution of sex differences in murine brain volume

Because of its unprecedented statistical power, our spatially fine-grained voxelwise analysis of normative sex differences in *MouseBase* simultaneously expands and refines current understanding of sex-biased brain organization in mice (**Fig 1**). Classical foci of sex-biased brain volume that were discovered by earlier histological analysis [BST, MEA, SDN-MPO^2,4,8,35^] emerge as the largest and most punctate effect size peaks of a much larger landscape of sex-biased regions - perhaps explaining why they were the first to be discovered histologically. Newly identified foci span cortical, hippocampal, thalamic, midbrain, pontine, medullary, and cerebellar territories. Several of these already have scattered prior reports of sex-biases at diverse non-volumetric levels of analysis^36–40^, and lie within circuits implicated in sex-biased domains of nociception^41,42^, auditory processing^43^, stress responses^44^, and social or reproductive behavior^45,46^. Such volumetric differences likely index underlying cellulo-molecular variation, as shown in our snRNAseq analysis of the cerebellum.

### Combined contributions of gonadal and sex chromosomal factors

Using parallel sMRI of wild-type and SCT mice, our study (i) successfully recapitulates the long-established causal role of gonadal hormones in shaping classical foci of sex-biased brain volume (BST, MEA, SDN-MPO and AVPV)^7,8,47^, and (ii) extends this knowledge by systematically quantifying the relative volumetric effects gonadal and sex-chromosomal factors across the newly expanded set of sex-biased brain regions we define in *MouseBase* (**Fig 2, Fig S2, Fig 3**).

First, we find that gonadal effects do not just co-localize with sex differences at previously described classical foci of predominantly male-sex biased volume, but also closely track the substantially expanded map of both male- and female-biased volumetric sex differences delineated in *MouseBase* (i.e. T>O where male>female and O>T where female>male, **Fig 2g,h, Fig S2c, Fig 3b**). Second, although gonadal effects are much more strongly coupled to regional sex differences than sex chromosome effects, we find that consideration of sex chromosomes alongside gonadal effects significantly improves prediction of volumetric sex differences - attesting to a combined contribution of these two causal factors. Specifically, consideration of sex chromosome effects improves prediction of volumetric sex differences within female-biased regions as a whole (**Fig S3a**), and at specific foci of both male- and female-biased regional brain volumes (**Fig S3b**), including regions classically considered to reflect purely gonadal effects (BST, MEA) as well as sex-biased regions of the cerebral cortex, cerebellar cortex and hippocampus. At some of the foci (BST, MEA, prelimbic cortex) gonadal and chromosomal effects appear to operate in opposition to one another - suggesting a “yin yang” interaction^48,49^ whereby two sex-biased factors can combine to lessen the eventual magnitude of a phenotypic sex differences **Fig 2g,h, Fig S2c, Fig 3b**). Third - and supporting their combinatorial contribution to sex differences we show that gonadal and sex chromosomal influences on cortical volume are in fact spatially coordinated with each other across the brain, and that the volumetric influence of sex chromosome complement reflects the contrast between XCD and YCD effects on volume. Although differences in XCD vs. YCD effects play a role in shaping sex differences, we find that XCD and YCD effects on volume show a strong positive correlation across the mouse brain - as they have previously been shown to in numerous sMRI studies of humans with sex chromosome aneuploidy^50–54^. The perhaps surprising observation of that highly heteromorphic X- and Y-chromosome can have convergent effects on biology has also been reported for autosomal gene expression in human cell lines^55^ - a phenomenon which we replicate herein across multiple murine cell types in our single cell analysis of cerebellar gene expression (**Fig 4**).

Convergence in XCD and YCD effects presumably reflects impacts of one or both of the two gene sets shared between and X- and Y-chromosome: pseudoautosomal region (PAR) genes (which show perfect sequence homology and still recombine between the X- and Y-chromosome, but are few in mice) and gametologs (which no longer recombine between X- and Y-chromosomes, but have nevertheless been conserved across species and retained high sequence homology between their X- and Y-linked copies). Comparative genetics and functional genomics of the sex chromosomes suggest that the gametologs may be especially important for convergent XCD and YCD effects^55–57^. At the same time, some gametologs appear to have developed partially functional divergence between their X- and Y-linked copies^57^, and these - together with many other genetic and epigenetic differences between the X- and Y-chromosome^58^ could explain those differences between XCD and YCD effects that appear to contribute to sex differences in regional brain volume (**Fig. 2, S3**).

### Relevance of findings for models of sex-biased brain organization in humans

Our findings provide important new data points for ongoing debates regarding the existence of sex-biased brain anatomy in humans^1^. The extensive sex differences in murine brain volume we report herein - often moderate to large in effect - arise without human cultural confounds and reflect the combined effects of gonadal and chromosomal sex factors that mice and humans share. Thus, unless human-specific factors (e.g. sociocultural influences) obscure gonadal and chromosomal effects on brain volume, our murine findings would predict the existence of regional neuroanatomical sex differences in humans. This theoretical prediction aligns with recent sMRI evidence for reproducible effects of both sex^1,14^ and sex chromosome dosage ^50–54,59^ on human brain anatomy.

It is notable that the brain regions we find to show volumetric effects of sex, gonadal and chromosomal effects in mice often lie within brain circuits that underpin domains of sex-biased murine behavior - raising the idea that volumetric sex differences may be a marker for cellulo-molecular sex differences that could contribute to behaviour. This potential linkage between volumetric and cellulo-molecular sex differences has been hinted at by prior work in mice^60,61^, is supported by our snRNAseq findings in the murine cerebellum (**Fig. 4**), and may also operate in humans ^62^ - suggesting a potential logic through which the spatial patterning of sex-biased brain anatomy in humans could inform our understanding of sex differences in the prevalence and presentation of so many brain-based medical conditions^1^.

For those brain regions that show convergent volumetric sex differences in humans and mice^14^, the mechanistic bases we discover in mice (**Fig. 4**) can be transferred to guide theory and research in sex differences in the human brain. For example, based on our findings in mice, we hypothesize that the female bias in volume of human frontal and sensorimotor cortices, and the male bias in human amygdala volume^1^ - which are both also seen in mice (**Fig. 1**) - reflects a mix of gonadal and sex chromosomal effects (**Fig. 3**).

### Limitations

The current study and its findings should be considered in light of several caveats and limitations. First, we focus on regional measures of brain volume, and future work will need to examine the many other properties of brain organization that can be queried by murine MRI (e.g. white matter organization). Second, MRI-based analysis of brain anatomy has both strengths and weaknesses relative to histologically based analysis methods. Our study harnesses these advantages, but does not avoid the weaknesses, such as limited information regarding those cellular and molecular features that presumably underlie volume variation (which we probe instead by snRNAseq). Third, our study is cross-sectional in adulthood, so does not speak to potential age-related variation in sex-biased brain anatomy or age-specific effects of gonads or sex chromosomes. Fourth, we do not have information on interindividual variation on hypothalamo-pituitary-gonadal axis functioning, but factors such as estrous cycle phase in females have been shown to modulate murine brain organization^63^ and may therefore modulate observed sex differences.

### Conclusions

Notwithstanding the above limitations and caveats, our study provides a new reference map of sex differences in murine brain anatomy and importantly annotates this map systematically for the causal influences of gonadal type and sex chromosome dosage. We dramatically expand the set of regions known to show sex-biased brain volume in mice, and localize these differences at a fine-grained voxel scale. This expanded map of sex differences encompasses over half of the mouse brain and includes distributed regions of both male- and female-biased volume - many of which are key nodes within well-described brain circuits subserving sex-specific reproductive and social behaviors. By integrating these maps from wild-type mice with coregistered MRI data from transgenic mice, we comprehensively characterize the relative contribution of gonadal type and sex chromosome complement to sex-differences in brain volume. These analyses recapitulate classical findings from experimental studies at large effect-size foci of male-biased volume, and newly show that gonadal influences are indeed the major contributor to regional sex differences in volume throughout the much-expanded set of sex-biased regions detected by MRI. However, we also find substantial evidence for sex chromosomal contributions to regional volumetric sex differences. This combination of gonadal and sex chromosomal contributions to sex differences is also evident at the level of cell-type specific gene expression profiles within regions of sex-biased anatomy. Because sMRI-derived maps of regional volumetric sex differences in the mouse brain can be compared with similar maps for the human brain, our study builds a bridge towards more refined mechanistic models for sex-biased brain development in humans.

## Methods

### Acquisition and preprocessing of murine neuroimaging data

*MouseBase* images were obtained as part of our study on autism and related neurodevelopmental disorders^21^. In that study 135 mouse models relevant to neurodevelopmental disorders were included; here we retained MRI scans of wild-type controls from 29 mouse models which met our inclusion criteria of having at least 5 male and 5 female wild-types, resulting in a total of 670 scans. SCT mouse MRI scans were acquired separately but using identical protocols as detailed below.

Sex Chromosome Trisomy (SCT) mice were originally obtained from Paul Burgoyne’s lab at the Crick Institute, and maintained at UCLA on an MF1 outbred background but with a uniform X chromosome identical in all mice (originally created by breeding an XO mother with an XY son). The Y chromosome in all mice originates from the 129 strain. SCT mice were maintained in using two separate pairings of (1) XXY-mothers with XY129 fathers to generate further XXY-mice and (2) XX mothers with XY-Sry fathers to generate four core genotype offspring. Full SCT breeding to generate all 8 possible genotypes was via crossing XXY-mothers with XY-Sry fathers. Brains were harvested using the same protocol as in *MouseBase* and shipped to Toronto for MRI imaging. In brief, MF1 mice bred at UCLA, using the Sex Chromosome Trisomy cross (**Fig 2a**) were perfused with 4% paraformaldehye and ProHance. Skulls were imaged at University of Toronto. Genotypes were determined by a combination of PCR to detect the Y^−^ chromosome and *Sry* transgene^64^, and interphase Fluorescent In Situ Hybridization to detect the number of X and Y chromosomes (Kreatek kit KI-30505, Leica Biosysystems, USA). Mouse numbers for imaging are given below.

### Mousebase data acquisition

Readers are referred to Ellegood et al^21^ for a detailed description of acquisition and processing methods. Briefly, all data were acquired ex-vivo on mouse brains kept within the skull after cardiac perfusion with fixative and gadolinium contrast agent. MRI scans were acquired 16 at a time^65^ in custom built solenoid coils on a 7T Varian (Agilent) MRI system or 8 at a time in similar coils in a 7T Bruker MRI. We used a T2-weighted, 3D fast spin-echo sequence, with a cylindrical acquisition of k-space, and with a TR of 350 ms, and TEs of 12 ms per echo for 6 echoes, field-of-view of 20 x 20 x 25 mm^3^ and matrix size = 504 x 504 x 630 giving an image with 40 µm isotropic voxels^66^.

### Image preprocessing

Anatomy measures were then derived using an image registration pipeline implemented in *PydPiper*^22^. We used a two-level framework, with the first level being a within-mouse model aligning all mutants and controls, and the second level across mouse models aligning average images produced in the first level together into a common space. Imaging alignment at each level was iterative; first, images were aligned using linear (6-parameter fit followed by a 12-parameter fit) and nonlinear registration. Following registration, we resampled the images using the appropriate transform and subsequently created a population average to represent the mean anatomy of the entire study sample. All registrations were performed using a combination of mni_autoreg tools^67^ and advanced normalization tools (ANTs)^68,69^. At the end of this process we have a transform taking each mouse MRI scan into the common space representing the average of all MRI scans. These transformations were then inverted and Jacobian determinants calculated and used as the measure of volume change at every voxel. As mentioned above, only data from 670 wild-type control mice from the 29 studies with sufficient number of male and female controls were retained to form *MouseBase*, while all 181 SCT mouse brain scans were retained.

### Mapping volumetric sex differences in *MouseBase*

We generated our map of normative sex differences in the brain using a linear mixed effects model, fitted using the lme4 package^70^ in R v4.3.2, computed separately at every voxel of the brain using RMINC (https://github.com/Mouse-Imaging-Centre/RMINC). The linear mixed effects model used was

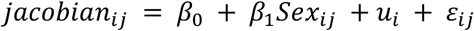

For the jth mouse in the ith mouse model, where

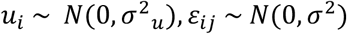

Equivalently, in lme4 notation:
jacobian ∼ Sex + (1|mouseModel)

Estimating a random intercept per mouse model has the effect of statistically removing effects that are due to the mouse model, which equally accounts for differences in originating vivarium, mouse background, and age.

The Jacobian determinant used in these models derives from the non-linear registration and thus represent volume differences after accounting for overall brain size [since global difference in brain size are accounted for by the linear (3 separate scales computed for the X, Y, and Z dimension) part of the transform]. Multiple comparisons were accounted for using the False Discovery Rate (FDR)^71^; degrees of freedom were estimated using the Satterthwaite approximation^72^. Cohen’s d effect sizes were computed from the marginal t-statistics using the following approximation:

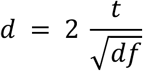

### Mapping gonadal, sex chromosome complement, X- and Y-chromosome dosage effects in SCT mice

The SCT mouse model was parameterized with the following terms:

| Shorthand | TvO | XYvXX | XCD | YCD | n |
| --- | --- | --- | --- | --- | --- |
| XX | Ovaries | XX | 2 | 0 | 20 |
| XXSry | Testes | XX | 2 | 0 | 32 |
| XY | Ovaries | XY | 1 | 1 | 26 |
| XYSry | Testes | XY | 1 | 1 | 19 |
| XXY | Ovaries | - | 2 | 1 | 21 |
| XXYSry | Testes | - | 2 | 1 | 17 |
| XYY | Ovaries | - | 1 | 2 | 24 |
| X YYSry | Testes | - | 1 | 2 | 22 |

We fitted two separate linear models:

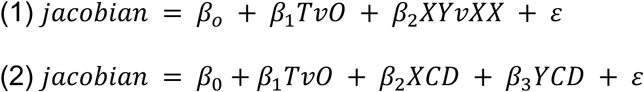

The first model only included mice with diploid sex chromosomes (i.e. the first four rows in the table above); the second model included all SCT mice. T-statistics were converted to effect sizes using the same approximation as above 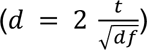, and effect sizes retained for XY vs XX chromosomes (XYvXX) from model (1) and testes vs ovaries (TvO), X chromosome dosage (XCD), and Y chromosome dosage (YCD) from model (2).

### Interrelating sex, gonadal, sex chromosome complement, X- and Y-chromosome influences in regions of sex-biased volume

We next sought to model how well the SCT terms (TvO, XYvXX, XCD, and YCD) predict and explain normative sex differences (MvF). All analyses were restricted to the SBM. We emphasized explanatory power over highest accuracy, and thus opted to use penalized regression splines implemented via general additive models (GAM) in the *mgcv* package in R^73^.

We tested eight different GAM models, in each case predicting the MvF effect sizes using combinations of SCT terms. The models where:

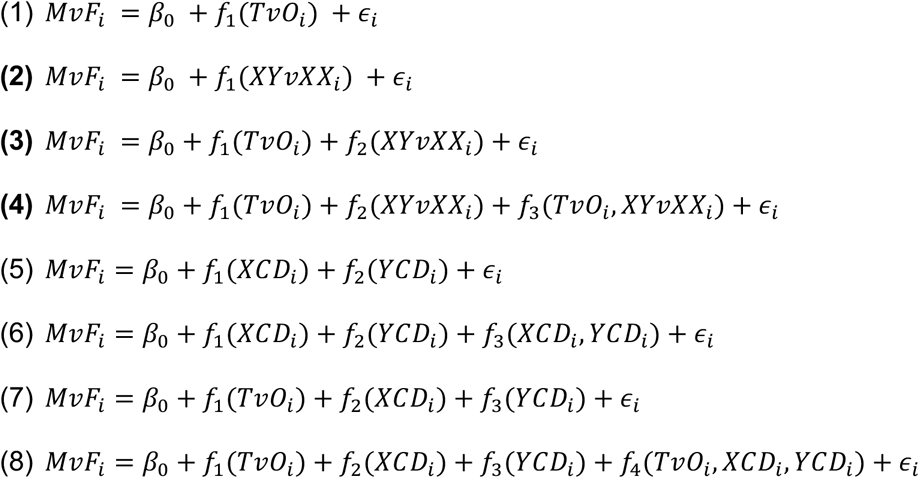

Where the functions *f* are mgcv’s *ti* or *tensor product interaction* functions, and ϵ_i_ ∼ *N*(0, σ^2^). Each *f* was heuristically constrained to have a maximum of 6 degrees of freedom, again operating on the principle that explanatory ability was valued over pure fitting accuracy. Model 8 was, for example, written as follows in R:

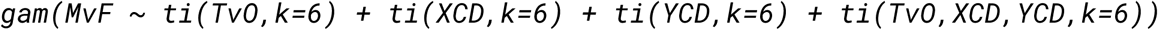

To estimate relative model performance while avoiding overfitting biases to the more complex models we followed a 10 fold cross-validation as well as permutation approach. In brief, we split the normative sex dataset of 670 MRI scans into equal splits of 335 MRIs each, fit the 8 GAM models on one half, then estimated R^2^ on the other half. We repeated this process 10 times. The SCT data stayed constant for each split. For each split we recreated the marginal t-statistics MvF map. Note that for computational tractability we estimated MvF as a linear, rather than a linear mixed effects, model, covarying for study rather than including study as a random intercept. The marginal t-statistics were converted to approximated effect size maps, again as above, and then used to compute the 8 GAM models. To evaluate the probability of the R^2^ values occurring by chance we estimated 100 permutations, using the same 50/50 splitting strategy as above, but this time sampling (without replacement) the sex assignment. Each of the 10-fold cross validation splits was then evaluated against each of the 100 permuted splits, with the number of times that the permuted splits R^2^ exceeded the CV splits R^2^ taken as the p-value. Finally, for **Fig. S2** we computed Pearson’s correlations between all the terms of the SCT dataset as well as normative MvF. Two additional SCT terms were included here, involving the algebraic addition or subtraction of the effect size maps of the two chromosome dosage terms.

### Single nucleus RNAsequencing

#### Tissue dissection, library preparation and sequencing

We focused snRNAseq analyses on a cerebellar region - the paramedian lobule (PRM) - which was a novel area of female-biased volume (**Fig. 1, S1**) that also displayed combined effects of both gonadal type and sex chromosome complement on volume (**Fig. 2**). This region was dissected from 24 mice encompassing 3 individuals from each of the 8 SCT model groups (mean age 65 days, range 60-70, no difference in age by genotype F=0.7, p=0.7). Dissection was conducted using the Franklin and Paxinos atlas^74^ as a reference. Breeding and genotyping of SCT mice was as described above under neuroimaging Methods. Original assigned genotypes based on interphase FISH were assessed based on *Xist* and *Uty* gene expression post-sequencing. We identified 2 mis-genotyped samples, leading to a final group number as follows: XXSry-(n = 3), XXSry+ (n = 4), XXY-Sry-(n = 3), XXY-Sry+ (n = 2), XY-Sry-(n = 3), XY-Sry+ (n =3), XY-Y-Sry-(n = 3), XY-Y-Sry+ (n = 3). Nuclei were isolated from cerebellar samples using the Nuclei EZ Prep Kit following the manufacturer’s experimental procedure (Sigma, St. Louis, MO, USA). Single-nuclei suspensions were then used to generate GEMs and libraries following the 10x 3′ single-cell RNA-seq V3.1 protocol (10x Genomics, Pleasanton, California, USA). The concentration of the libraries was determined using the Qubit Fluorometric Quantitation method (Thermofisher Scientific, Waltham, MA, USA), and quality was assessed using the Agilent TapeStation system (Agilent, Santa Clara, CA, USA). The pooled libraries were then sequenced on the Novaseq S4 2×100 (Illumina, San Diego, CA, USA) in the UCLA Broad Stem Cell Research Center at ∼25k reads/cell. ***Preprocessing and quality control.*** Raw base call (BCL) files were processed and individually aligned to the *mus musculus* genome assembly GRCm38 (mm10) using CellRanger version 6.0.2 (10X Genomics). Intronic reads were included as snRNA-seq profiles primarily nuclear transcripts. Sample Fastq files were processed using CellBender version 0.3.0 to remove signals derived from technical artifacts such as ambient RNA molecule contamination^75^. Sample-wise digital gene expression matrices, in which each row represents read counts for a gene and each column an individual cell, were loaded, combined, and subsequently analyzed using the R package Seurat version 4.1.1. Nuclei with <200 or >7,500 UMIs, feature counts < 500 or > 35,000, mitochondrial gene expression >3%, and hemoglobin gene expression > 5% were filtered out as poor quality or contaminated cells or potential doublets. DoubletFinder version 2.0.4 was used to further identify potential contaminating doublet cells which were removed from the final dataset^76^.

#### Clustering cell and identifying cerebellar cell types

The gene expression matrix was processed using the standard Seurat workflow, including data normalization, identification of variable genes, data scaling, and principal component analysis. To correct for batch effects driven by differences in sequencing date, cells were integrated via Harmony. Single cells were projected onto two dimensions using Uniform Manifold Approximation Projection (UMAP) and assigned to clusters using Louvain clustering. To identify the cell type identity of each cluster, cluster-wise expression of canonical cell type markers derived from literature of commonly found cell types in the brain were assessed (including reference cerebellar data from^28^). Clusters that showed unique expression of known marker genes were used as evidence for cell type annotation. Annotations were further validated using the Allen Institute’s MapMyCells tool, which accepts a gene expression matrix to assign cell type identities from Allen Institute taxonomies. Cell type proportions within genotype groups were calculated and significant differences were evaluated using ANOVA.

#### Computing sex differences in gene expression per cell type

Genes expressed in at least 15% of cells were selected for differential gene expression analysis. The limma R package version 3.58.1 was used to identify differentially expressed genes (DEGs) between XY_Testes and XX_Ovaries SCT groups with “Sequencing Date” included as a covariate, allowing for determination of male-biased and female-biased sex differences in gene expression for each cell type cluster. Multiple testing was corrected for with the Benjamini-Hochberg method. DEGs with adjusted p-value < 0.05 and with at least a 0.1 log2 fold change in gene expression were identified as significant and included in downstream pathway and logFC correlation and regression analyses.

#### Computing gonadal, sex chromosome complement, X- and Y-chromosome dosage effects on cell type expression

The same workflow as described above was used to assess changes in gene expression caused by individual sex-biasing factors by utilizing the same linear model used to analyze the anatomical data, which incorporated gonad type (TvO; “Gonad” in **Fig. 4e**), sex chromosome complement (XYvXX: “SexChrom” in **Fig. 4e**)), X-chromosome dosage (2 vs 1: “XCD” in **Fig. 4e**), and Y-chromosome dosage (linear increase from 0 to 1 to 2: “YCD”, in **Fig. 4e**), with Sequencing Date added as a covariate. Multiple testing was corrected for with the Benjamini-Hochberg method. DEGs with adjusted p-value < 0.05 and with at least a 0.1 log2 fold change in gene expression were identified as significant and included in downstream pathway and logFC correlation and regression analyses.

#### Relating sex effects on gene expression to effects of gonadal type, sex chromosome complement, X- and Y-chromosome dosage

Sex differences in gene expression were related to gonadal and sex chromosomal effects (i.e. TvO, XYvXX, XCD and YCD) by first extracting logFC values for all terms in each cell type, and then computing pairwise Pearson correlations between all terms per cell type - separately for autosomal and X-chromosome genes (all pairwise correlations shown in **Fig. S4f**). Linear models were also fitted to estimate regression slopes for TvO, XYvXX, XCD and YCD terms in estimation of sex effects on gene expression in each cell type (separately for autosomal and X-chromosome genes). These correlations and regressions were conducted for the set of genes that a significant sex effect on expression in at least one cell type (and excluding XIST and TSIX in calculations for X-chromosome genes since the extreme expression changes for these genes would dominate estimation of correlation and regression coefficients). For each cell type, correlation and regression coefficients for TvO, XYvXX, XCD and YCD terms in estimation of sex effects on expression were combined to visualize the degree to which expression effects of each term recapitulated the observed sex effect (**Fig. 4e**).

#### Pathway analysis

Significant female and male-biased DEGs were used as input to test enrichment of pathways per cell type from the GO Biological Processes (GO:BP) and GO Cellular Components (GO:CC) databases, which was assessed with Fisher’s exact test followed by multiple testing correction with the Benjamini-Hochberg method. Pathways at adjusted p-value < 0.05 were deemed significantly overrepresented.

## Acknowledgements

This work was supported by the NIH-NICHD grant R01HD100298 (A.P.A, A.M.G, J.L, A.R, J.S, X.Y), NIH-NIMH grant R21MH129020 (X.Y); the Intramural Research Program of the National Institute of Mental Health (NIMH) (National Institutes of Health [NIH] Annual Report Number: ZIAMH002949, A.R.); the NIH Training Grant in Genomic Analysis and Interpretation T32HG002536 (J.S). The authors thank the University of California, Los Angeles Technology Center for Genomics and Bioinformatics for sequencing of the snRNA-seq data.The authors would also thank Xuqi Chen for assistance.

## Data availability

Original MRI data is available, in part, at https://www.braincode.ca/content/public-data-releases#dr001. Remaining data will be released very soon at the same location alongside publication of ^21^. Key statistical maps are available at https://osf.io/4cbas. All source code to generate all results and figures is available at https://git.fmrib.ox.ac.uk/jason/sct-to-parse-sex-differences.

---

**Fig S1.**
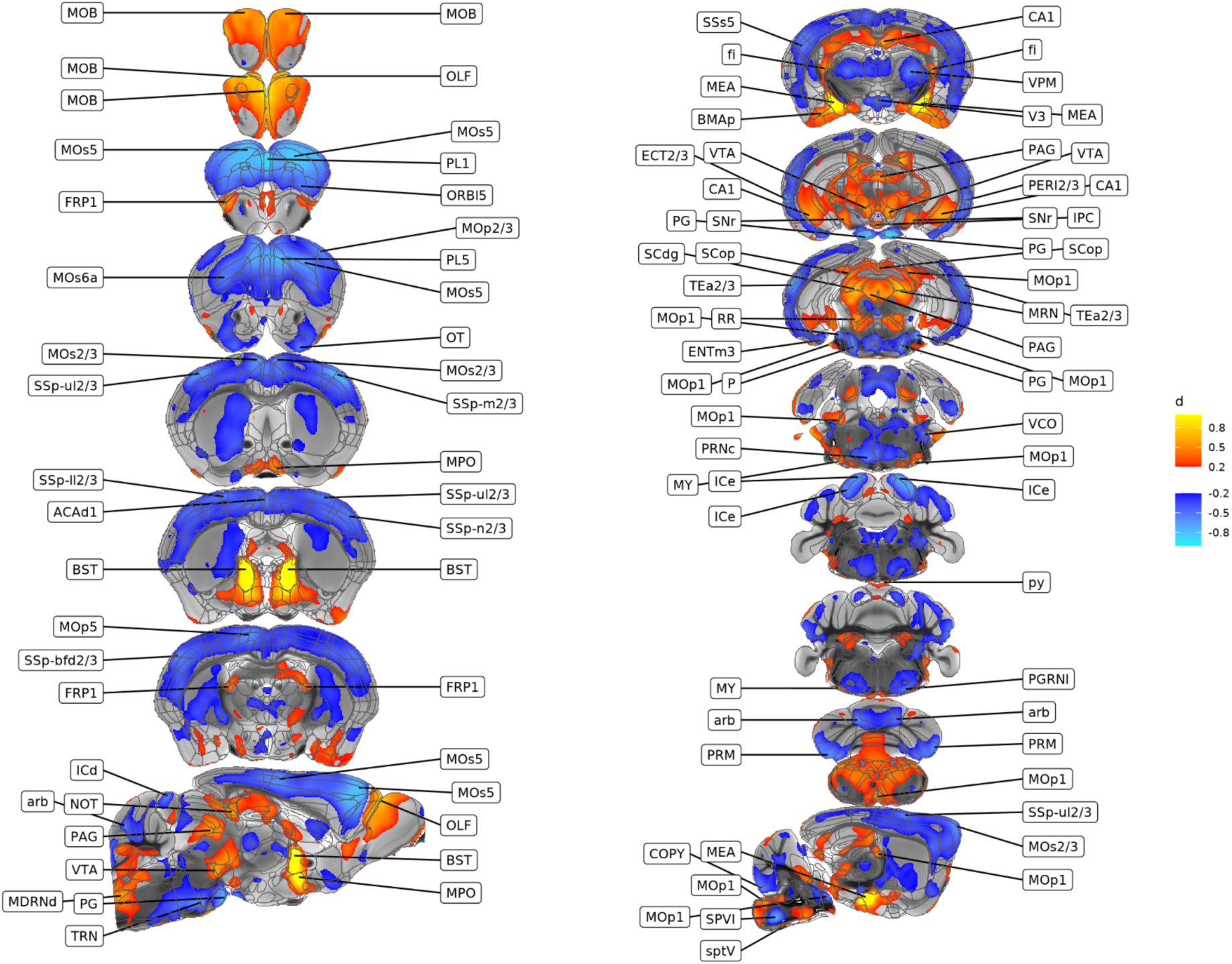
Regions of statistically significant sex differences in murine brain volume color coded by effect size. Fine coronal slices and selected sagittal slices through the mouse brain show regions of statistically significantly sex-biased volume surviving FDR correction for multiple comparisons at q<0.05 (these are finer slices of the map from Fig 1d). Colors encode effect sizes for the observed volumetric sex difference (female> male - blue; male>female - red). **Abbreviations:** ACAd1: Anterior cingulate area, dorsal part, layer 1; BMAp: Basomedial amygdalar nucleus, posterior part; BST: Bed nuclei of the stria terminalis; CA1: Field CA1; COPY: Copula pyramidis; ECT2/3: Ectorhinal area and Layer 2 and 3; ENTm3: Entorhinal area, medial part, dorsal zone, layer 3; FRP1: Frontal pole, layer 1; ICd: Inferior colliculus, dorsal nucleus; ICe: Inferior colliculus, external nucleus; IPC: Interpeduncular nucleus, caudal; MDRNd: Medullary reticular nucleus, dorsal part; MEA: Medial amygdalar nucleus; MOB: Main olfactory bulb; MOp1: Primary motor area, Layer 1; MOp2/3: Primary motor area, Layer 2 and 3; MOp5: Primary motor area, Layer 5; MOs2/3: Secondary motor area, layer 2 and 3; MOs5: Secondary motor area, layer 5; MOs6a: Secondary motor area, layer 6a; MPO: Medial preoptic area; MRN: Midbrain reticular nucleus; MY: Medulla-other; NOT: Nucleus of the optic tract; OLF: Olfactory areas-other; ORBl5: Orbital area, lateral part, layer 5; OT: Olfactory tubercle; P: Pons-other; PAG: Periaqueductal gray-other; PERI2/3: Perirhinal area, layer 2 and 3; PG: Pontine gray; PGRNl: Paragigantocellular reticular nucleus, lateral part; PL1: Prelimbic area, layer 1; PL5: Prelimbic area, layer 5; PRM: Paramedian lobule; PRNc: Pontine reticular nucleus, caudal part; RR: Midbrain reticular nucleus, retrorubral area; SCdg: Superior colliculus, motor related, deep gray layer; SCop: Superior colliculus, optic layer; SNr: Substantia nigra, reticular part; SPVI: Spinal nucleus of the trigeminal, interpolar part; SSp-bfd2/3: Primary somatosensory area, barrel field, layer 2 and 3; SSp-ll2/3: Primary somatosensory area, lower limb, layer 2 and 3; SSp-m2/3: Primary somatosensory area, mouth, layer 2 and 3; SSp-n2/3: Primary somatosensory area, nose, layer 2 and 3; SSp-ul2/3: Primary somatosensory area, upper limb, layer 2 and 3; SSs5: Supplemental somatosensory area, layer 5; TEa2/3: Temporal association areas, layer 2 and 3; TRN: Tegmental reticular nucleus; V3: third ventricle; VCO: Ventral cochlear nucleus; VPM: Ventral posteromedial nucleus of the thalamus; VTA: Ventral tegmental area; arb: arbor vitae; fi: fimbria; py: pyramid; sptV: spinal tract of the trigeminal nerve.

**Fig S2.**
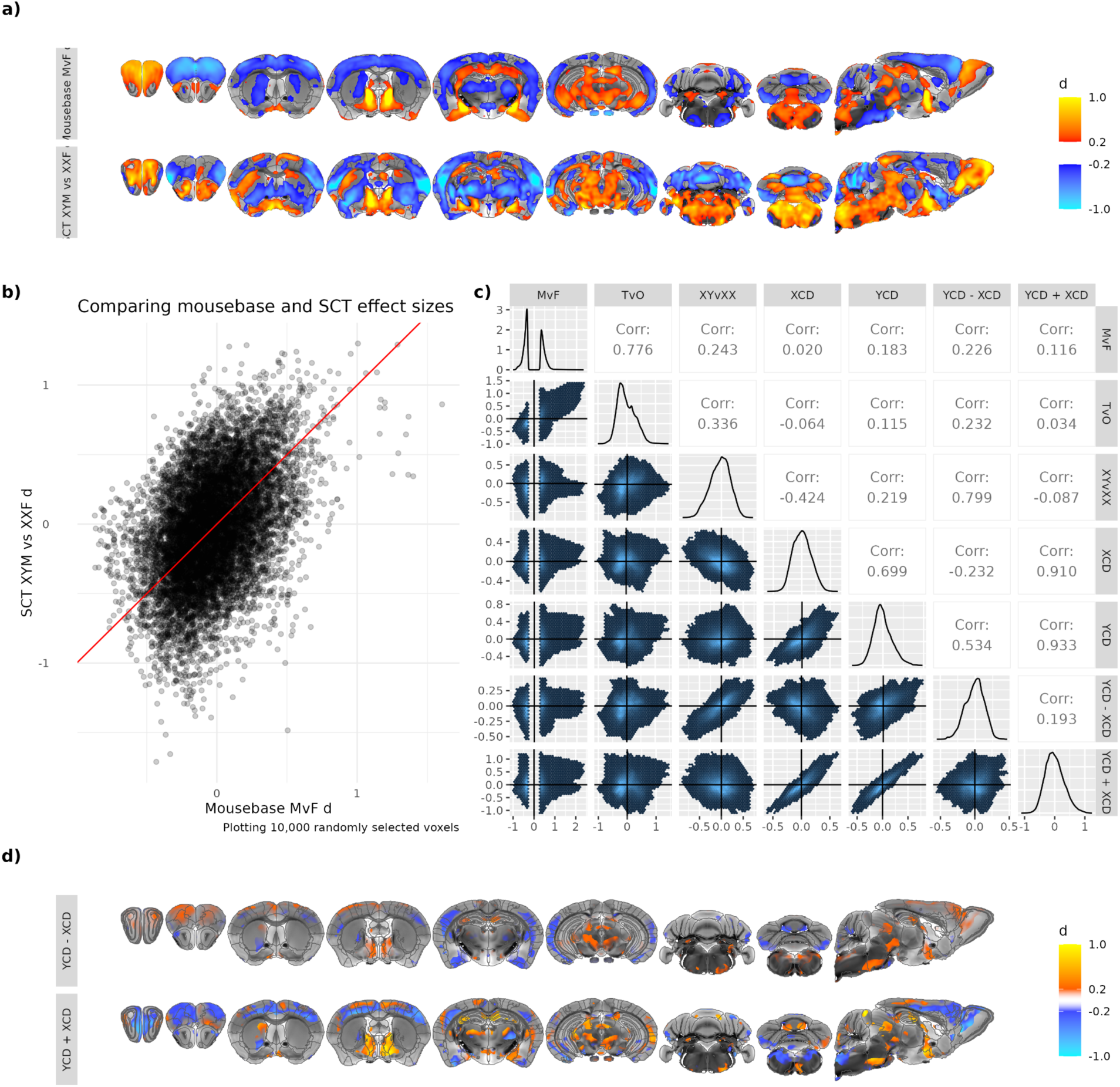
Spatial correlations between observed volumetric sex differences in *MouseBase* and SCT cohorts, and amongst volumetric effects of gonadal type, sex chromosome complement, and dosage of X- and Y-chromosomes (XCD, YCD) as estimated in the SCT cohort. **a**, Selected coronal and sagittal views of the mouse brain showing voxel-wise volumetric sex differences within *MouseBase* (top row - wild type male vs. female contrast for all voxels in the sex-biased mask (SBM)) and the SCT model (bottom row - XY-testes vs. XX-ovaries group contrast for all voxels with |d|≥0.2). Voxels are color coded by the direction of volumetric sex differences (red - male-biased; blue - female-biased). **b,** Scatter plot showing the correlation between voxel-wise effects of sex in *MouseBase* and SCT datasets (i.e. maps in panel a) for 10k randomly selected voxels within the *MouseBase* SBM. The red line indicates the identity line. **c,** Scatterplot matrix showing cross-voxel correlations amongst the magnitude (standardized effect size) of normative volumetric sex differences within the *MouseBase* SBM and the magnitude of volumetric effects for gonadal type (TvO), sex chromosome complement (XYvXX) and dosage of X and Y chromosomes (XCD and YCD, respectively) as estimated within the SCT dataset (i.e. maps from Fig. 2b). We also include correlations for the **d**, algebraically derived difference between YCD and XCD effects (YCD-XCD, top row) and their sum (YCD+XCD, bottom row).

**Fig S3.**
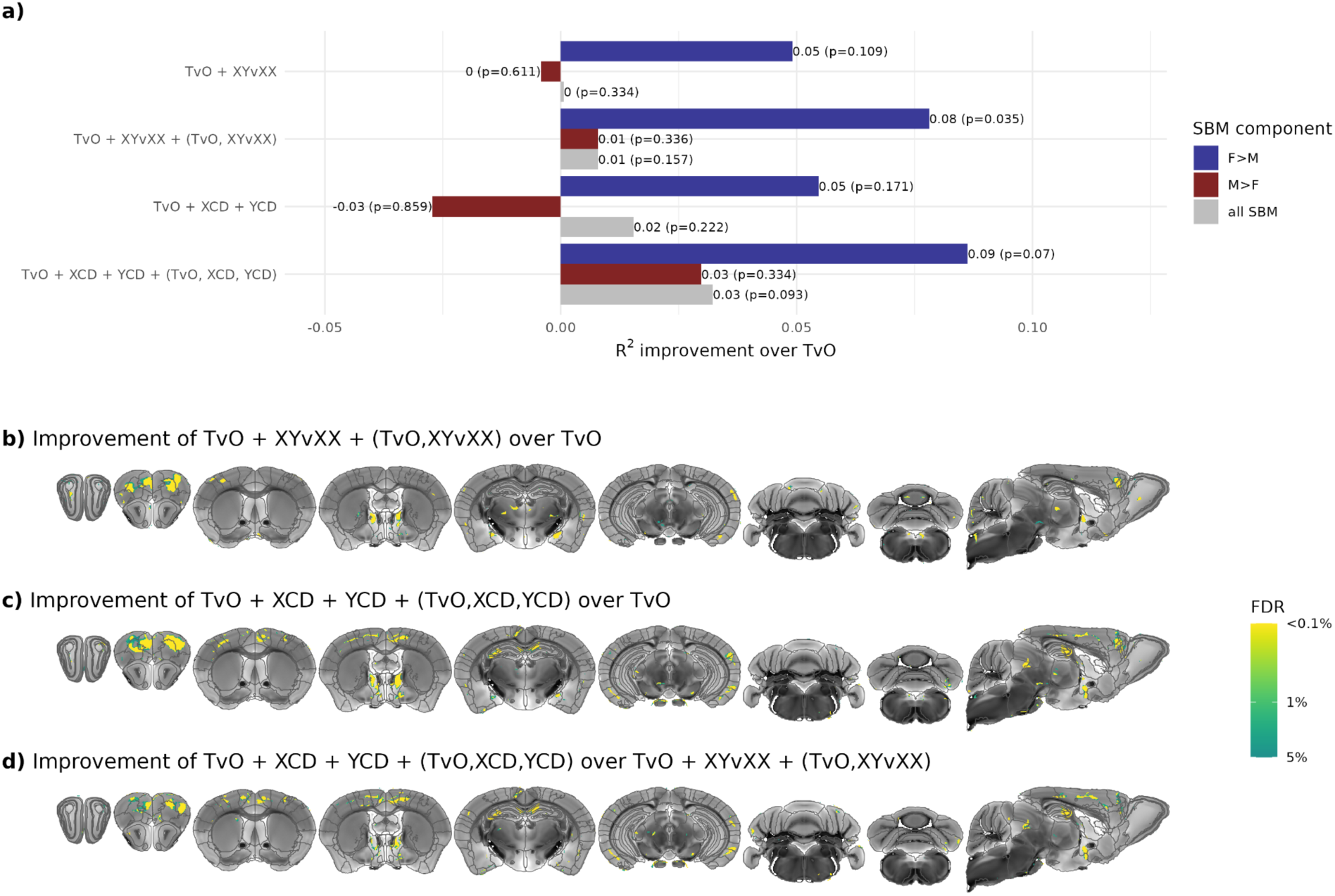
Incremental predictive power of adding sex chromosome terms to a gonadal term in prediction of volumetric sex differences. **a.** The magnitude and statistical significance of R^2^ changes for prediction of sex differences within the SBM from addition of chromosomal terms (combinations of XYvXX, XCD, YCD terms and their interactions) compared to a model with just a gonadal (TvO) term (blue - female-biased SBM (F>M); red - male-biased SBM (M>F); gray - whole SBM). E.g. addition of XYvXX effects and their interaction with TvO effects (TvO, XYvXX) to a base model with just TvO effects explains 8% more spatial variance in the magnitude of normative sex differences within female biased SBM voxels (blue colored bar) **b,c.** Voxel wise maps within the SBM showing regions where addition of sex chromosome terms yields a statistically significant improvement in the prediction of volumetric sex differences (reduction in root mean squared error) compared to a model with just a gonadal (TvO) term. **b** shows that areas of the frontal cortex, BNST, and MeA improve by including the base sex chromosome terms. **c** shows further increase in the area of the cortex improved by also considering sex chromosome dosage terms; hippocampal regions also show an effect here. **d** shows regions where the model in d) improves prediction on sex differences over the model in c).

**Fig S4.**
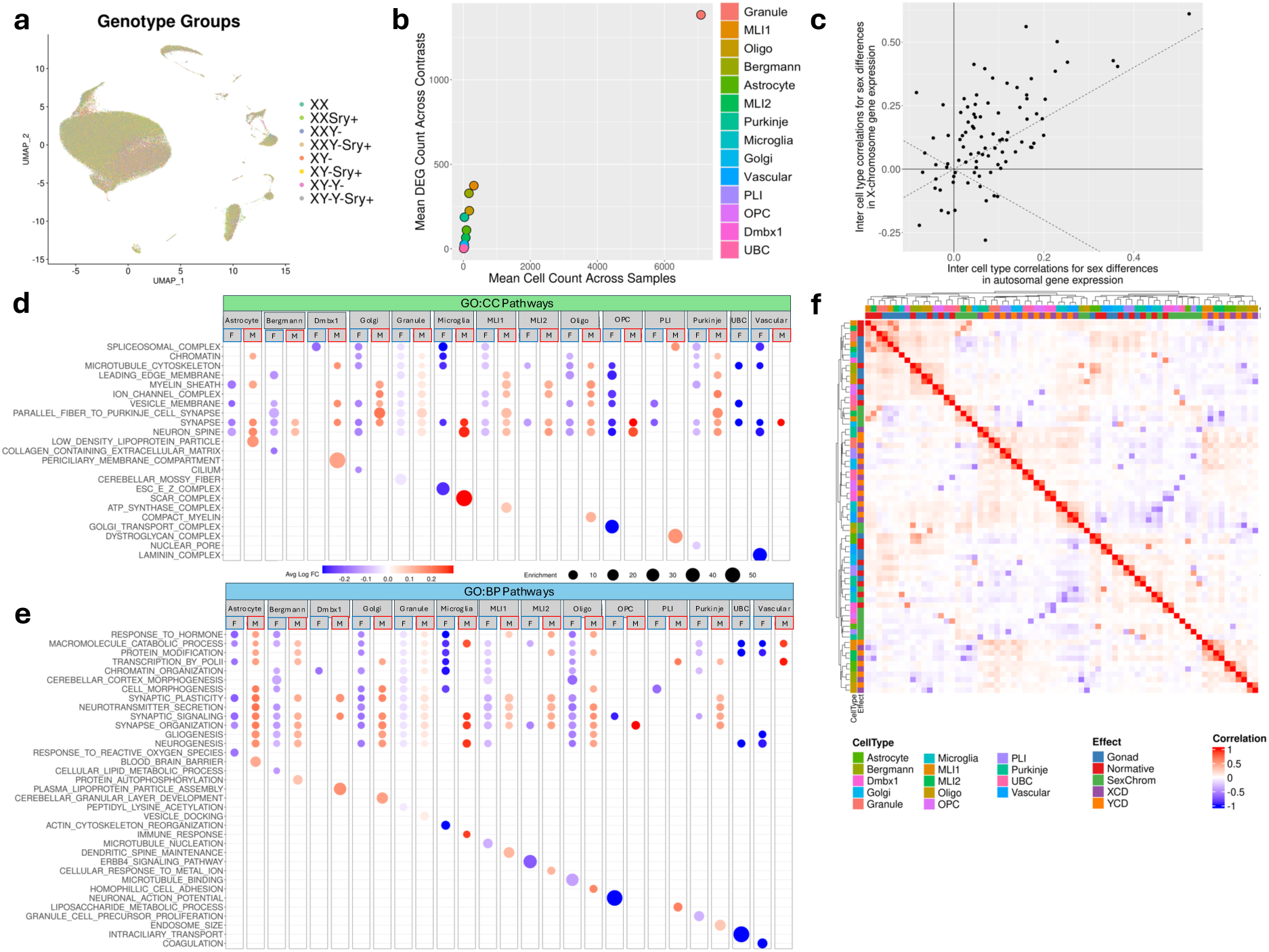
Supplementary analyses and results for cerebellar snRNAseq analyses. **a.** UMAP showing clustering of 8 genotype groups of the SCT mouse model **b**. Dotplot showing the mean cell count by mean DEG count for all cell types across samples. **c**. Dotplot showing inter-cell type similarity in fold change correlation coefficients between autosomal and X-chromosome genes. Xist and Tsix excluded as outliers to trend. **d-e.** Top enriched pathways identified from sex-biased DEGs across cerebellar cell types, based on (**d**) Gene ontology cellular component (GO:CC) and (**e**) Gene ontology biological processes (GO:BP) databases. Dots are colored by average fold change between sexes (XX_Ovaries vs XY_Testes for female-biased (blue); XY_Testes vs XX_Ovaries for male-biased (red)) for significant DEGs overlapping with indicated pathway in specific cell type. Dot size is proportional to enrichment score. **f**. Correlation heatmap of logFC values across cerebellar cell types and sex effects based on Pearson correlation coefficients calculated pairwise across cell type x sex effect DEG sets. Rows and columns annotated by cell type and sex effect. Cells colored by strength and direction of correlation. Black squares highlight positive cross cell-type correlations between XCD and YCD effects on gene expression.

## Notes

### Competing Interest Statement

The authors have declared no competing interest.

